# Transcriptomic Analysis Identifies Candidate Genes for Differential Expression during *Xenopus laevis* Inner Ear Development

**DOI:** 10.1101/2023.12.29.573599

**Authors:** Selene M. Virk, Casilda Trujillo-Provencio, Elba E. Serrano

## Abstract

**Background:** The genes involved in inner ear development and maintenance of the adult organ have yet to be fully characterized. Previous genetic analysis has emphasized the early development that gives rise to the otic vesicle. This study aimed to bridge the knowledge gap and identify candidate genes that are expressed as the auditory and vestibular sensory organs continue to grow and develop until the systems reach postmetamorphic maturity.

**Methods:** *Affymetrix* microarrays were used to assess inner ear transcriptome profiles from three *Xenopus laevis* developmental ages where all eight endorgans comprise mechanosensory hair cells: larval stages 50 and 56, and the post-metamorphic juvenile. Pairwise comparisons were made between the three developmental stages and the resulting differentially expressed *X*. *laevis* Probe Set IDs (Xl-PSIDs) were assigned to four groups based on differential expression patterns. DAVID analysis was undertaken to impart functional annotation to the differentially regulated Xl-PSIDs.

**Results:** Analysis identified 1510 candidate genes for differential gene expression in one or more pairwise comparison. Annotated genes not previously associated with inner ear development emerged from this analysis, as well as annotated genes with established inner ear function, such as *oncomodulin*, *neurod1,* and *sp7*. Notably, 36% of differentially expressed Xl-PSIDs were unannotated.

**Conclusions:** Results draw attention to the complex gene regulatory patterns that characterize *Xenopus* inner ear development, and underscore the need for improved annotation of the *X. laevis* genome. Outcomes can be utilized to select candidate inner ear genes for functional analysis, and to promote *Xenopus* as a model organism for biomedical studies of hearing and balance.

## Introduction

The coordinated activity of multiple genes underlies many complex biological processes including organ development, cell differentiation, and the onset and persistence of diseases. High-throughput methods, such as microarrays and RNA-Seq, can quantify the expression levels of thousands of transcripts simultaneously, permitting a system-level analysis of transcriptomes. The use of global profiling technologies enables the discovery of expression profiles that correlate with biological processes, which can result in the generation of hypotheses regarding the genes involved in the maintenance of the phenotype in question (McDermott et al. 2007; Friis et al. 2011; Darville and Sokolowski, 2013; Gu et al., 2016; Baxi et al, 2023; Tisi et al, 2023).

The inner ear is an example of an intricate organ system whose development requires the controlled expression of manifold genes, many of which have yet to be fully defined (Alsina et al, 2009; Wu et al., 2012; Mackowetzky et al., 2021). The inner ear’s role as a sensory receptor for mechanical forces imparts its vital function as an organ essential for vertebrate survival and reproduction. The start of the current millennium ushered in the genomics era and subsequently, various genes that are essential for inner ear development and function have emerged. For example, evidence suggests that a number of transcription factors such as *DLX5*, *SIX1*, and *GATA2* are associated with hearing loss and/or vestibular dysfunction and many Online Mendelian Inheritance in Man^®^ genes have been linked to deafness and vestibular disorders (Merlo et al., 2002; Zheng et al., 2003; Haugas et al., 2010; Ramirez-Gordillo et al.. 2015; Taiber et al, 2022). However, in comparison with other organ systems such as the eye, heart, and kidney, far less is known about the inner ear’s developmental transcriptome, and the control of genetic networks that typify inner ear development (Giraldez and Fritzsch, 2007; Broto et al., 2021; Mackowetzky et al, 2021). Investigations that seek to understand how the expression profiles of functionally relevant genes change during inner ear development and maturation, are necessitated by the pressing need to develop therapeutic interventions for disorders of hearing and balance that affect almost half a billion humans (Agrawal et al., 2009; Eshraghi et al., 2013; Lustig and Akil, 2019; Tabier et al., 2022). It is anticipated that transcriptomic profiling will uncover crucial inner ear genes whose function is relevant for prevention or repair of malfunctioning auditory and vestibular systems, as well as for *in vitro* studies of inner ear organs in stem cell derived systems (Koehler and Hashino, 2014; Lee and Waldhaus, 2022).

The inner ear’s unique role, sequestered location within the temporal bone, and relatively minute size make studies with human tissue impractical. Consequently, investigators who study hearing and balance rely on a spectrum of animal models and advocate for the advantages of their particular organism as a model for investigating inner ear structure and function (Giraldez and Fritzsch, 2007; Mackowetzky et al, 2021; Lee and Waldhaus, 2022). Common justifications for species selection include the organism’s relevance as a mammalian model (rodents, cats, primates), its regenerative potential (fish, amphibians, birds), or its suitability for transgenesis (*Danio Rerio*, *Xenopus*, *Mus musculus*). In fact, foundational understanding of the physiological role of the inner ear’s mechanoreceptor hair cells was pioneered in non-mammalian models through seminal biophysical investigations of amphibian and turtle hair cells (Hudspeth et al., 1977; Ricci et al., 2002; Kozlov et al. 2007).

Research presented here implements the amphibian, *Xenopus*, for developmental investigations of inner ear transcriptomics using microarray approaches. A popular and important genus for studies of embryogenesis (Gurdon, 2014; Kostiuk and Khoka, 2021), *Xenopus* has been relatively underutilized for studies of organogenesis, neural systems, or aging, especially in comparison with zebrafish, an organism that is evolutionarily far more distant from humans than amphibians (Nakatani et al., 2007;). With two sequenced genomes (*X. laevis* and *X. tropicalis*), Resource Centers that can produce and stock transgenic animals (Vize and Zorn, 2017; Horb et al, 2019) and documented neuroregenerative potential (Burns et al., 2013; Phipps et al., 2020), *Xenopus* is poised to contribute to genomic investigations of neural and sensory processes (Nenni et al., 2019; Exner and Willsey, 2021). Previous transcriptomics studies featuring *Xenopus* have profiled multiple organs and developmental stages, but few have focused on the inner ear and its development (Baldessari et al., 2005; Powers et al., 2012; Langlois and Martyniuk 2013). The paucity of global profiling studies that feature the inner ear represents a missed opportunity to identify new genes for inner ear and organ function. Technologies that measure transcript abundance across multiple genes also afford the opportunity to discover groups of genes with synchronized changes in expression. Identification of such genes has the potential to uncover previously unknown functional relationships.

We implemented a microarray approach for analysis of the *Xenopus* inner ear transcriptome at three developmental ages that represent anatomically distinct stages of inner ear organ formation: *s50*-s52 (*s50*), *s56*-s58 (*s56*), and juvenile (aged 3 months). Larval age *s50* represents the youngest developmental age when all auditory and vestibular end organs have been anatomically characterized as fully formed (*s50*) (Bever et al., 2003; Quick et al., 2005). The *s56* age was selected as an intermediate larval age during which the inner ear undergoes an expansive growth period as evidenced by both the development of new hair cells and innervation by the eight cranial nerve (Díaz et al., 1995; Lopez-Anaya et al., 1997; Bever et al., 2003). Finally, the juvenile age was selected to identify alterations in gene expression related to the structural and functional changes that occur after metamorphosis, when *Xenopus* enters reproductive age. For example, although animal size increases after metamorphosis, very few hair cells are added to the sensory field and the 8^th^ nerve undergoes extensive myelination as compared with larval ages (Diaz et al, 1995; Lopez-Anaya et al., 1997).

Differential expression analysis of pairwise comparisons between developmental stages enabled identification of transcripts with the most variable expression levels between ages, as well as those that were predominantly expressed during a single age. The genes identified from the pairwise comparisons were subjected to functional annotation clustering using the Database for Annotation, Visualization and Integrated Discovery (DAVID); transcripts upregulated in the juvenile in comparison to the two larval stages were categorized as the cohort with the greatest enrichment score. The analysis presented here emphasized identifying the genes with the greatest expression changes to produce a robust data set that expands the characterization of three distinct developmental stages of the *Xenopus* inner ear. Research outcomes enhance the annotation of the *Xenopus* genome, and offer a resource that can be used by the inner ear development community to generate hypotheses about sensory organ formation, as well as for investigators interested in drug discovery (Wheeler and Liu, 2012; Liu and Yang, 2022;Cousins, 2022), and the use of stem cells for investigations of inner ear organ systems (Zhang and Hu, 2012; Roccio, 2021).

## Results

### Evaluation of replicate data for technical consistency of sample and array methods

Raw, log_2_ transformed, data for each of the three developmental stages and the associated replicates are shown in the boxplots in Fig. 1A. Visual inspection of boxplots prior to preprocessing showed similar medians in stage replicates, with the *s56* replicates appearing the least variable. In order to achieve the broadest intensity distribution, raw array data were preprocessed using GCRMA. Preprocessing resulted in normalized data with reduced variability in comparison to raw data (Fig. 1B). Examination of scatter plots of all samples plotted against one another (Fig. 1C), showed the greatest correlation to be between replicate samples and the least between different developmental stages, with the exception of the *s50* replicates. Histograms of all developmental stages were also constructed. The histograms displayed similar intensity distributions with the greatest intensity peak at the lowest observed intensity level (Fig. 1C), demonstrating the sensitivity of the GCRMA method to low intensity levels in comparison to other preprocessing methods such as RMA (Wu et al., 2004). It was concluded that the stringent surgical procedures and RNA isolation methods utilized in this study (Trujillo-Provencio et al., 2009) enhanced reproducibility and minimized technical experimental variation, based on (1) the similarities observed in both the replicate raw and preprocessed data, and (2) the lower observed variability between replicates than between developmental stages.

**Fig. 1.**
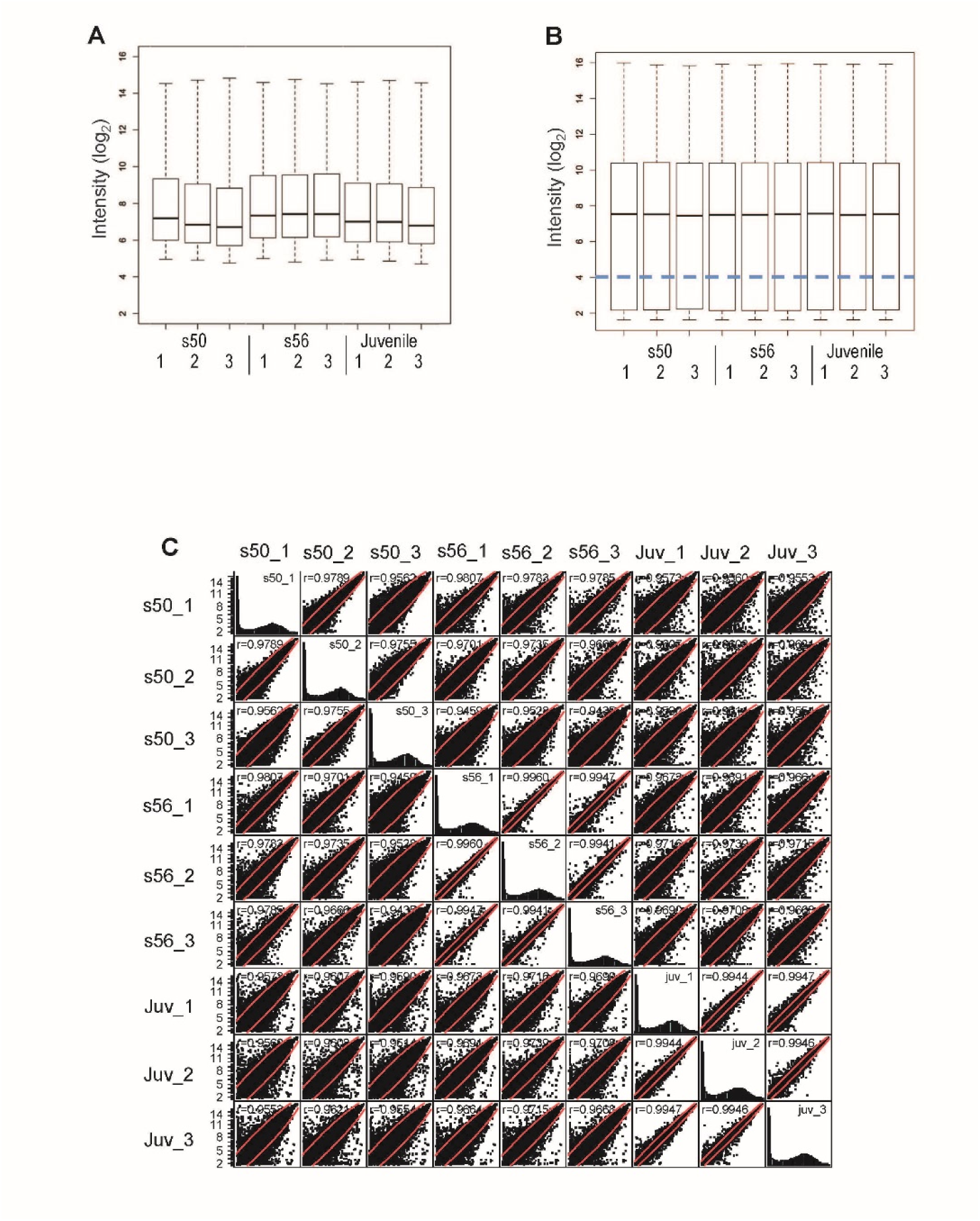
Comparison of microarray data distributions for samples of different ages. **(A, B, C)** Box plots of Affymetrix Genome 2.0 array data before **(A)** and after preprocessing with GCRMA **(B)**. (**C**) Scatterplots of the Xl-PSID intensity values from the three pairwise comparisons show greatest similarity between replicate comparisons of the same animal age. Fluorescence intensity values are in arbitrary units of fluorescence (a.u.).

### Differential expression analysis identified 1510 differentially expressed Xl-PSIDs (DEX) that were assigned to seven differential expression categories

The objective of this study was to determine gene expression changes during inner ear development using *Xenopus laevis* as a model. To achieve this objective, pairwise comparisons between the three developmental stages were made using the convention that a positive fold change specifies greater expression in the older stage. Data that met the filter criteria (q-value ≤ 0.01; fold change ≥ 1.5) were used for analysis. The filter criteria were implemented to focus on differences with a greater likelihood of biological significance, but to also include changes between genes that may be represented at low abundance such as transcription factors.

Volcano plots of the comparisons are shown in Fig. 2 A-C, with the dotted line crossing the x-axis at the significance level of q-value ≤ 0.01. The stages analyzed in this study included two different larval stages (*s50* and *s56*) and the metamorphosed juvenile (*Juv*). The fewest differentially expressed Xl-PSIDs (DEX) were found in the pairwise comparison featuring the two larval stages (Fig. 2A). The proportion of significant Xl-PSIDs increased in the *Juv-s56* comparison (Fig. 2B) and, finally in the *Juv-s50* plots (Fig. 2C), more data points were significant than in any other comparison. The heatmap shown in Fig. 3A displays hierarchical clustering and expression differences between all samples after filtering by statistical significance and fold change. In the heatmap, all replicates clustered by stage and the two larval stages clustered together.

**Fig. 2.**
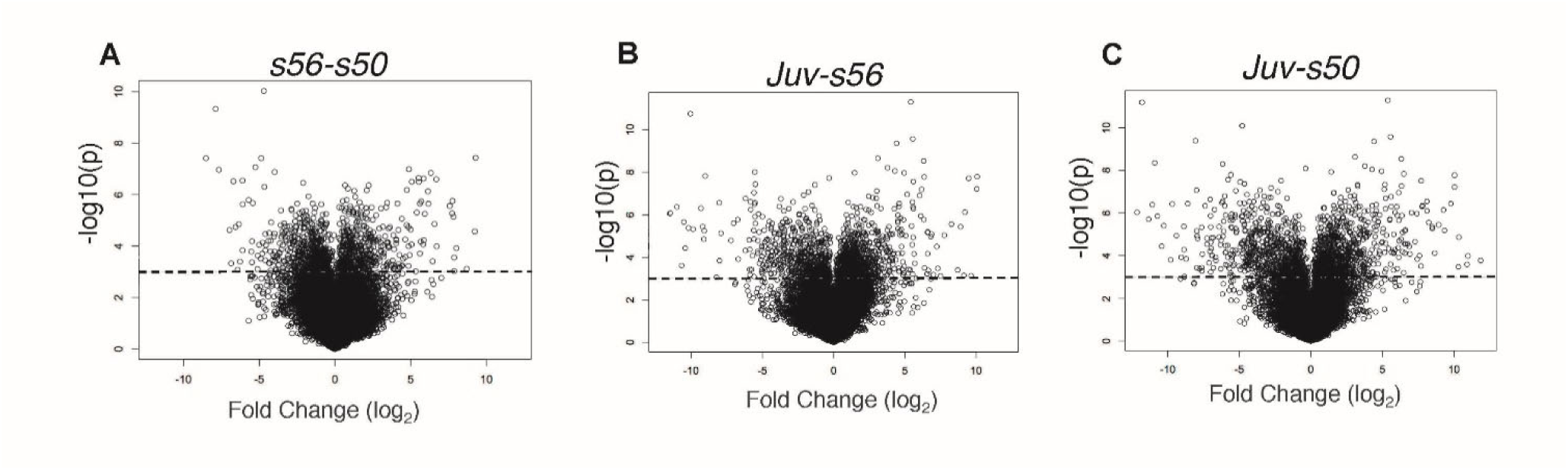
Results of pairwise comparisons between the ages. **(A,B,C)** Volcano plots of stage pairwise comparisons: s56-s50 **(A)**, Juv-s56 **(B)**, and Juv-s50 **(C)**. The Juv-s50 plot displays the greatest variation in the magnitudes of the log_2_ fold change in the differential analysis. Dotted line represents −log10(p) = 3.12; data points above this line are statistically significant at q ≤ 0.01.

**Fig 3.**
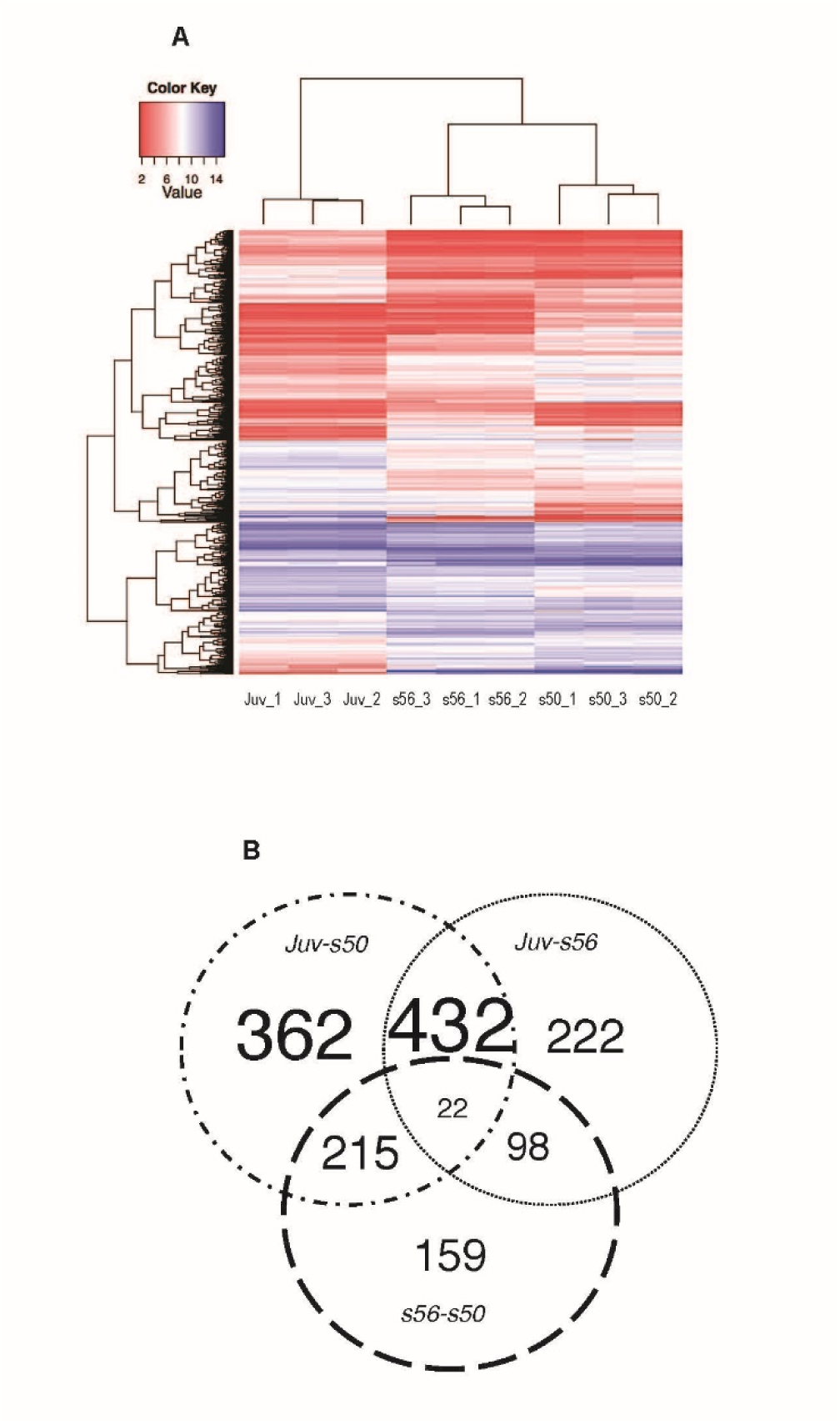
Heatmap and Venn Diagram of the 1510 differentially expressed Xl-PSIDs. **(A)** The heatmap shows the results of hierarchical clustering of all developmental stage replicates. Clear differences are visible between the developmental stages and the replicates from the two larval stages are clustered together. **(B)** Venn diagram depicting the distribution of Xl-PSIDs differentially expressed in one, two, or all three stage pairwise comparisons, with the greatest contribution to the total arising from Xl-PSIDs differentially expressed in the both the *Juv-s50* and *Juv-s56* comparisons. The threshold for differential expression was set at q-value ≤ 0.01 and minimum fold change difference of 1.5 a.u.

The resulting DEX could be assigned to seven categories based on their expression pattern (Fig. 3B). The division into the seven differential expression categories served as the overall focal point of the downstream analysis. The most populated categories comprised Xl-PSIDs that were differentially expressed in one (Group A) or two (Group B) pairwise comparisons, these two groups accounted for 98% of the DEX. The least populated category (Group C) encompassed Xl-PSIDs differentially expressed between all three stages, with 2% of the DEX total.

### Characteristics of Xl-PSIDs differentially expressed in a single pairwise comparison (Group A)

The greatest number of Xl-PSIDs in Group A were noted in the *Juv-s50* category, followed by the *Juv-s56* and *s56-s50* comparisons (Fig. 3B). In Group A, 34-38% of Xl-PSIDs differentially expressed in a single comparison were unannotated. The majority (61%) of the 159 DEX in the *s56-s50* comparison were upregulated in *s50* (*UP_s50*) and the observed fold changes ranged from −4.6 to 6.4 (Fig. 4A, Table 1). Among the most upregulated Xl-PSIDs in this comparison, two were unannotated and three were annotated with the following gene symbols: *atp6v1g3*, *LOC398308* (lectin type 2), and *tmod4*. Two Xl-PSIDs among the top expressed in *s56* also lacked annotation. The remaining four from the top *UP_s56* Xl-PSIDs corresponded to the gene symbols: *cldn1*, *slc25a13*, *klf9-b*, and *hhip*. The Xl-PSID annotated as *cldn1* (claudin 1), with a fold change of 4.9, was the Xl-PSID with the second highest fold change in this group (the highest fold change was for an Xl-PSID that was not annotated). The hedgehog interacting protein (*hhip*) is an inhibitor of hedgehog signaling, which is involved in anterioposterior patterning in *Xenopus* (Waldman et al., 2007).

**Fig. 4.**
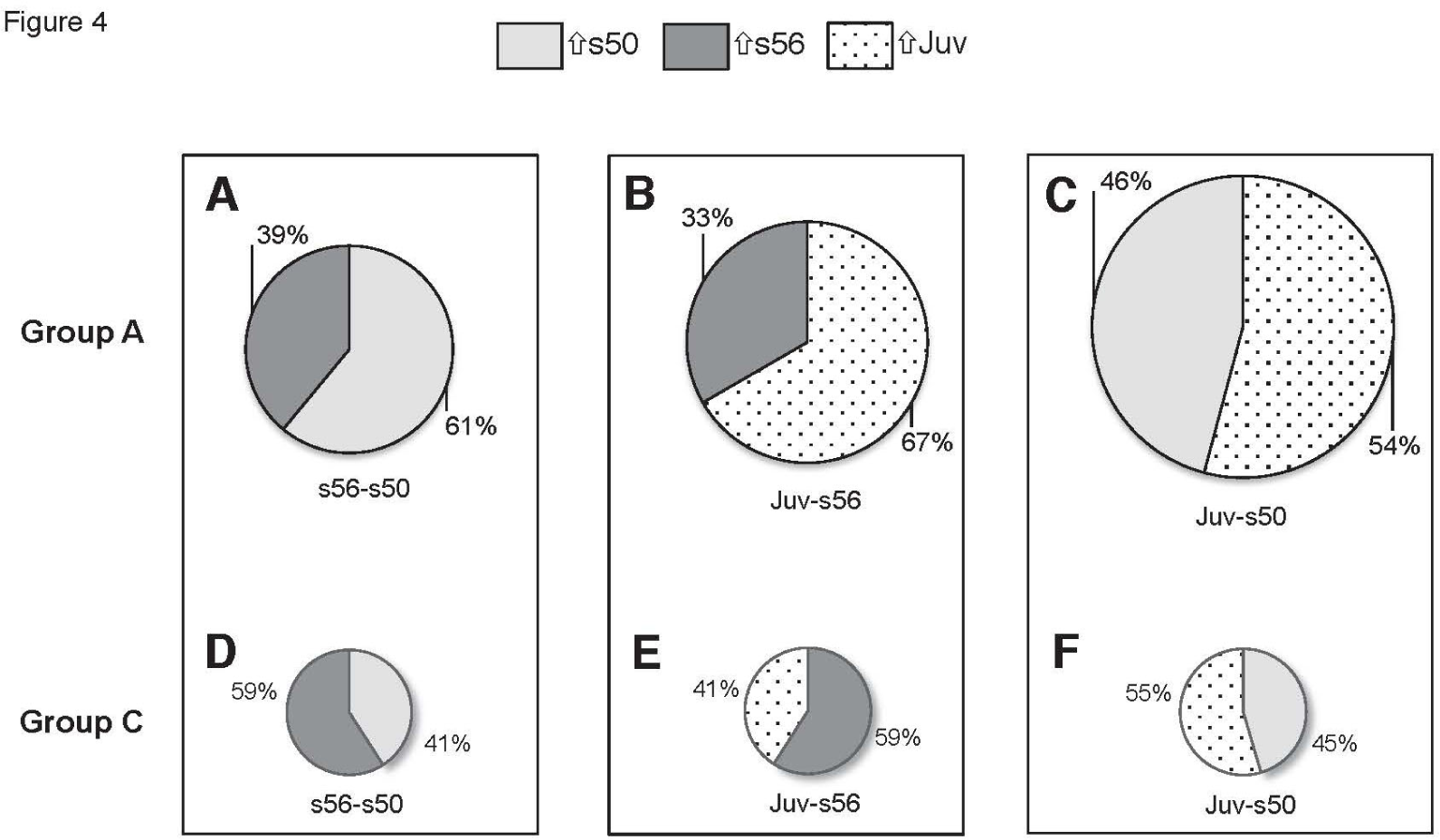
The smallest differential expression cohort was comprised of the 22 Xl-PSIDs differentially expressed in all three pairwise comparisons. **(A-C)** Microarray analysis uncovered 743 Xl-PSIDs that met the criteria for differential expression in only one pairwise comparison. In comparisons featuring the juvenile stage, the majority of Xl-PSIDs were upregulated in this stage as opposed to either of the larval stages. **(D-F)** The Xl-PSIDs differentially expressed in all comparisons had greater upregulation in s56 when compared to both the larval s50 and juvenile stages, in contrast to the pattern observed in Xl-PSIDs differentially expressed in only a single comparison. Circle size adjusted to reflect number of genes.

**Table 1.**
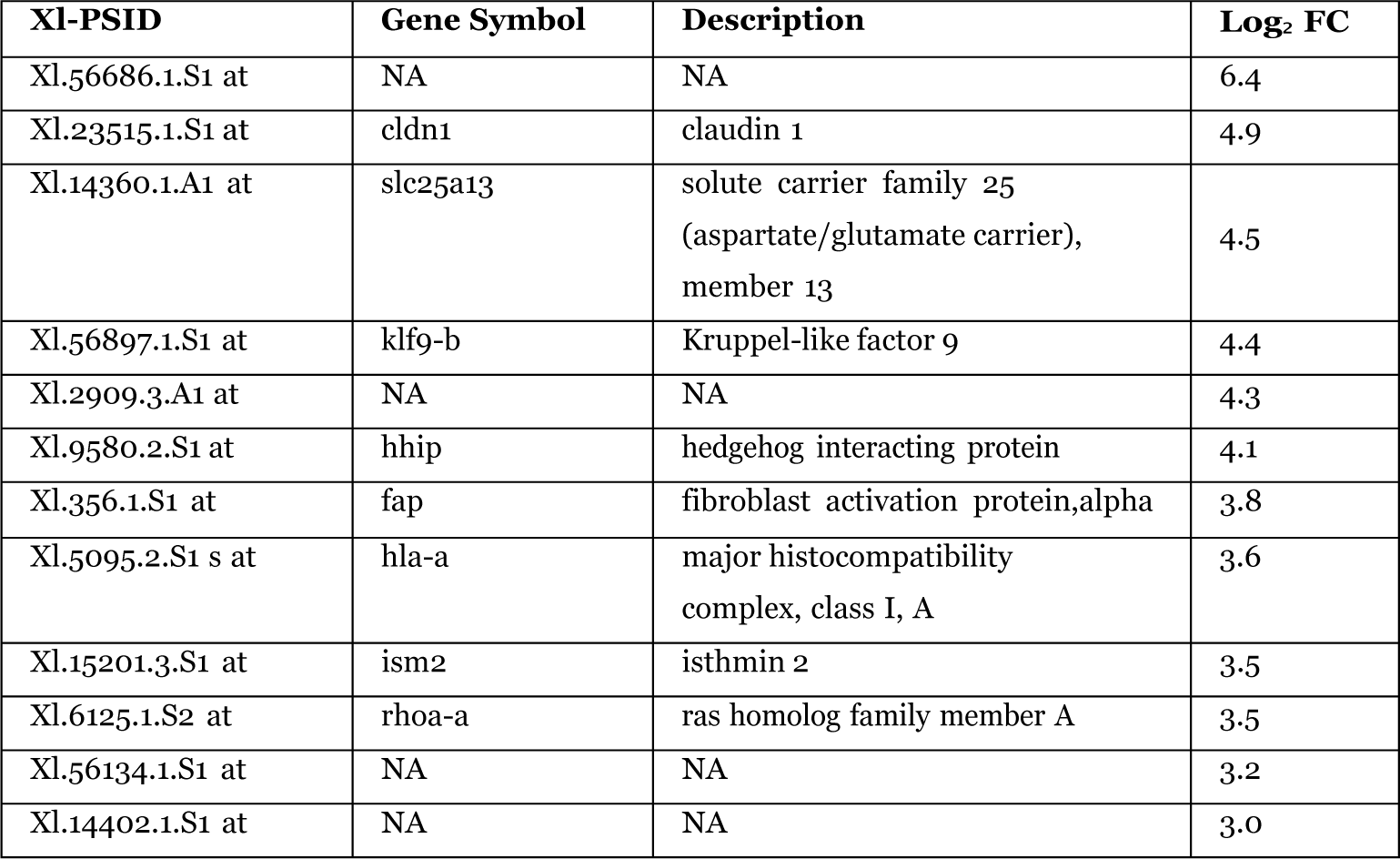
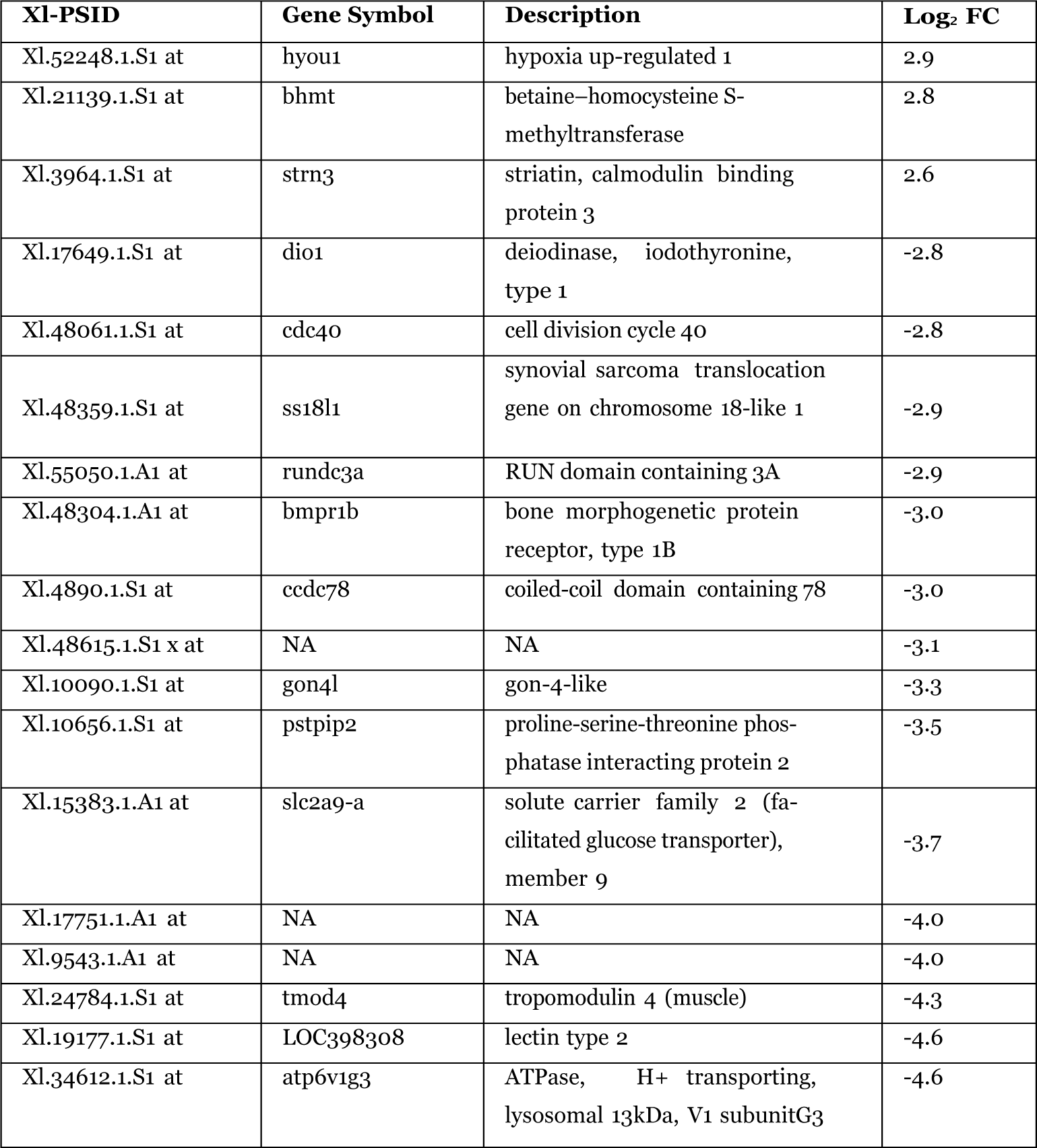
Table of the top 15 up/downregulated Xl-PSIDs differentially expressed in the s56-s50 comparison (Group A)

In the *Juv-s56* comparison, in contrast to the *s56-s50* comparison, greater upregulation (67%) was seen in the more mature developmental stage (Fig 4B). This comparison contained the second greatest count of DEX seen in this study and the fold change range was greater than in *s56-s50*, ranging from −10.7 to 9.6 (Table 2). The two Xl-PSIDs *UP_*56 with the greatest fold changes (−10.7 and −8) were annotated as mediator complex subunit 22 (*med22*) and major histocompatibility complex, class I, A (*hla-a*). The Xl-PSID annotated as deiodinase, iodothyronine, type 2 (*dio2*) had the fourth highest fold change of −5.8 and is involved in cochlea development in mice by way of local regulation of thyroid hormone (Campos-Barros et al., 2000). The Xl-PSID *UP_Juv* with the greatest fold change was identified as calpastatin-like (*calp1*), which is an inhibitor of a family of calcium activated proteases known as the calpains, which are involved in apoptotic processes (Rojas et al., 1999).

**Table 2.** Table of the top 15 up/downregulated Xl-PSIDs differentially expressed in the Juv-s56 comparison (Group A).

| **XI-PSID** | **Gene Symbol** | **Description** | **Log<sub>2</sub> FC** |
| --- | --- | --- | --- |
| Xl.503.2.S1 at | calp1 | calpastatin-like | 9.6 |
| Xl.630.3.A1 a at | cae-a | preprocaerulein | 7.0 |
| Xl.48479.1.S1 at | c11orf65 | Chromosome 11 open reading frame 65 | 6.7 |
| Xl.630.2.S1 s at | xt6l-a | XT-6 like precursor | 6.6 |
| Xl.33507.1.S1 a at | NA | NA | 6.6 |
| Xl.9135.1.A1 at | NA | NA | 6.4 |
| Xl.941.1.S1 at | NA | NA | 5.7 |
| Xl.5860.2.S1 at | ckm | creatine kinase, muscle | 5.4 |
| Xl.23640.1.S1 at | gng7-a | guanine nucleotide binding protein (G protein), gamma 7 | 5.3 |
| Xl.25551.2.S1 a at | NA | NA | 4.9 |
| Xl.51302.2.A1 at | NA | NA | 4.9 |
| Xl.194.1.S1 at | ptgds-b | prostaglandin D2 synthase<br>21kDa (brain) | 4.8 |
| Xl.37580.1.S1 at | mis18a | MIS18 kinetochore protein A | 4.6 |
| Xl.45948.2.S1 at | rpl36 | ribosomal protein L36 | 4.2 |
| Xl.48441.1.S1 at | hspb6 | heat shock protein, alpha-<br>crystallin-related, B6 | 4.2 |
| Xl.52756.3.S1 at | NA | NA | -3.6 |
| Xl.48824.1.S1 at | fam101b | family with sequence similar-<br>ity 101, member B | -3.7 |
| Xl.20526.1.S1 at | NA | NA | -3.7 |
| Xl.4366.1.A1 at | mki67 | marker of proliferation Ki-67 | -4.0 |
| Xl.1103.1.S1 at | LOC397864 | cytokeratin type I | -4.1 |
| Xl.2824.1.S1 at | NA | NA | -4.4 |
| Xl.56215.1.S1 at | NA | NA | -4.5 |
| Xl.5979.1.S1 at | NA | NA | -4.7 |
| Xl.54905.1.S1 at | NA | NA | -4.8 |
| Xl.26182.1.A1 at | NA | NA | -5.0 |
| Xl.53068.2.A1 at | NA | NA | -5.0 |
| Xl.21638.1.A1 at | dio2 | deiodinase, iodothyronine,<br>type 2 | -5.8 |
| Xl.17389.1.S1 at | olfm4 | olfactomedin 4 | -5.9 |
| Xl.30381.1.A1 at | med22 | mediator complex subunit 22 | -8.0 |
| Xl.5095.3.S1 a at | hla-a | major histocompatibility<br>complex, class I, A | -10.7 |

In contrast to both the *s56-s50* and *Juv-s56* comparisons, the *Juv-s50* comparison had a similar proportion of upregulated Xl-PSIDs between the older (54%) and younger ages (Fig 4C). In this comparison, the Xl-PSID fold-changes ranged from −9.8 to 11.9 (Table 3). The Xl-PSID most upregulated in *s50* was annotated as *pavlb.2* and the second most upregulated was oncomodulin (*ocm.2*), with a fold change of −9.1. Oncomodulin, also known as parvalbumin-β, is a member of the parvalbumin family of calcium binding proteins that are expressed in the mammalian cochlea and differentiates outer from inner hair cells (Sakaguchi et al., 1998). The expression level of oncomodulin decreases as development progresses in the rat inner ear (Yang et al., 2004), which was also observed in this analysis. One of the *UP_s50* Xl-PSIDs was transcription factor *neurod1-b*, with fold change of −6.6. The transcription factor neurogenic differentiation 1 (*neurod1*) is an important factor in the development of both the cochlea and the vestibular apparatus (Liu et al., 2000) and it represses the transformation of neuronal sensory cells into hair cells (Jahan et al., 2010). The most upregulated Xl-PSIDs in the juvenile stage in this comparison were the hemoglobins *hbg2-a* and *hbg1*, with fold changes of 11.9 and 10.3 respectively.

**Table 3.**
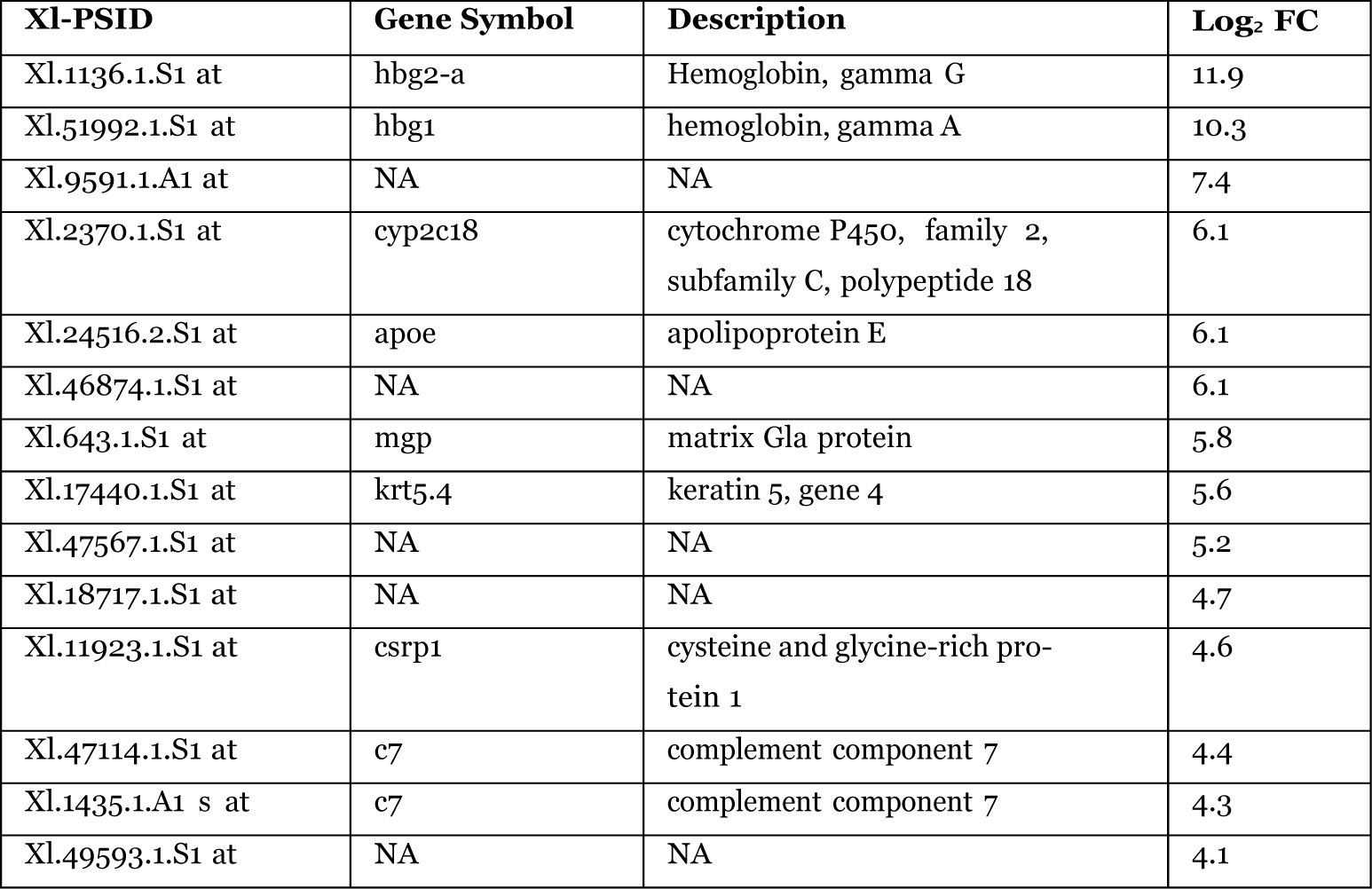
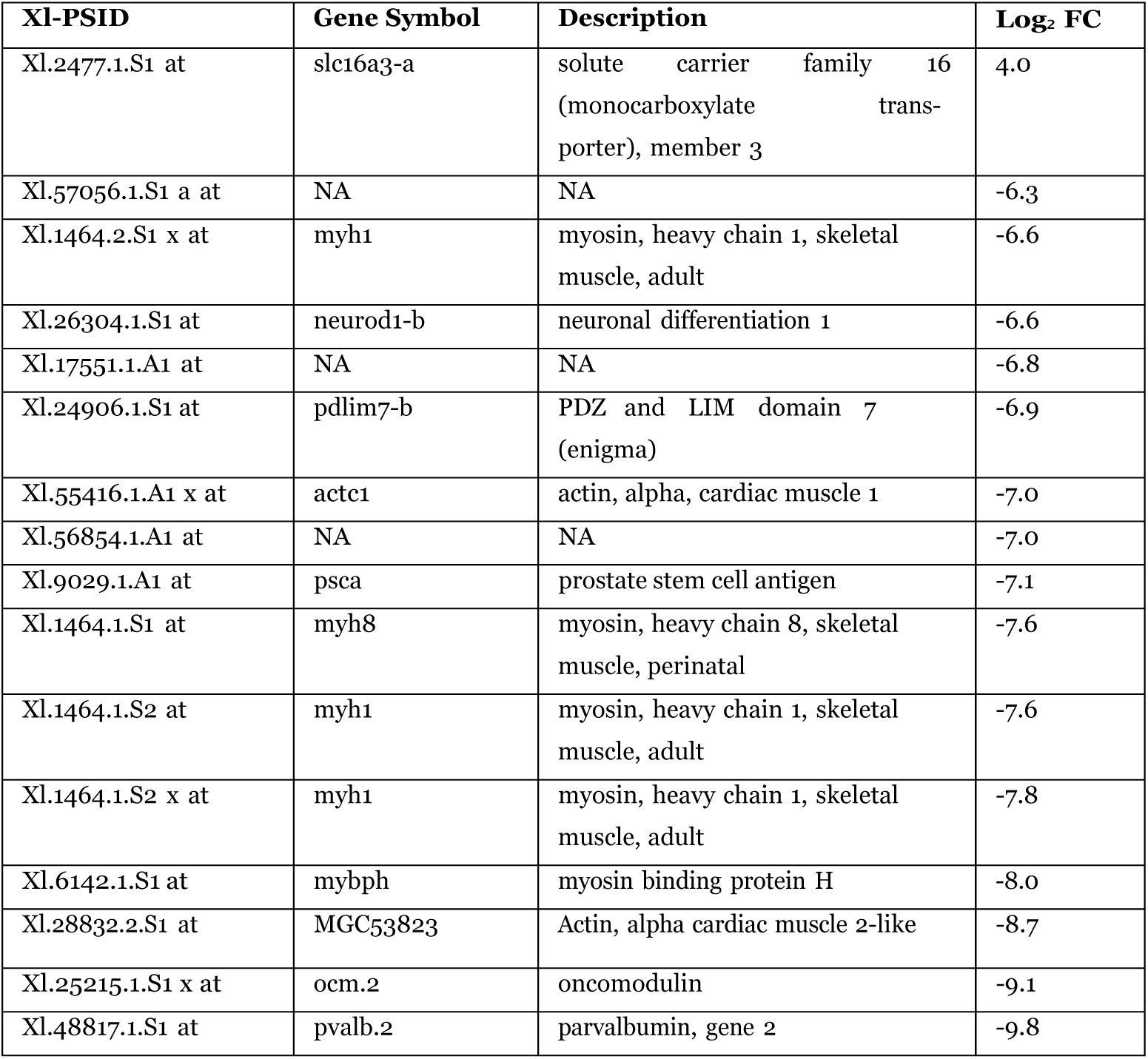
Table of the top 15 up/downregulated Xl-PSIDs differentially expressed in the Juv-s50 comparison (Group A).

### Analysis of Xl-PSIDs differentially expressed in two comparisons facilitated a “stage_centric” view of differential expression (Group B)

Xl-PSIDs that were differentially expressed in two comparisons (Group B) were termed *stage_centric*. Using the *s50* age as an example, the set of Xl-PSIDs differentially expressed when *s50* was compared to both *s56* and juvenile would represent the group of *s50_centric* Xl-PSIDs.

Analysis of DEX in *s56-s50* ∩ *Juv-s50* (*s50_centric*) provided information regarding differentially regulated genes in *s50* relative to a more developed larval stage and the completely metamorphosed juvenile. Genes that are *UP_s50* in this context could represent a cohort with decreasing expression during development. Most (55%) of the Xl-PSIDs in this category were upregulated in *s50* (Table 4). There were 215 DEX in *Juv-s50* ∩ *s56-s50*, 36% of which did not have a gene annotation. The Xl-PSID with the greatest upregulation in *s50* was annotated as dentin sialophosphoprotein (*dspp*), with fold changes of −8.9 (*Juv-s50*) and −6.3(*s56-s50*) [Table 5]. Mutations in *dspp* have been implicated in dentinogenesis imperfect, which can present with an autosomal dominant form of hearing loss (Xiao et al., 2001). Also among the most upregulated *s50* Xl-PSIDs were several genes with functional roles in cellular structure, two of which were the keratin genes *krt.5.6* and *krt14*, both with log_2_ fold changes of −8. The annotated Xl-PSID with the greatest downregulation in *s50* was annotated as bone gamma-carboxyglutamate [gla] protein (*bglap*), with fold changes of 8.5 and 7.8, compared to juvenile and *s56*, respectively.

**Table 4.** Xl-PSIDs differentially expressed in two pairwise comparisons (Group B) presents a stage centric view.

|  | **$s56-s50 \cap Juv-s50$** |  | **$s56-s50 \cap Juv-s56$** |  | **$Juv-s56 \cap Juv-s50$** |  |
| --- | --- | --- | --- | --- | --- | --- |
|  | Up<br>s50 | Down<br>s50 | Up<br>s56 | Down<br>s56 | Up<br>Juv | Down<br>Juv |
| XI-PSID<br>Counts | 119 | 96 | 48 | 50 | 208 | 224 |

**Table 5.** Table of the top 15 up/downregulated Xl-PSIDs differentially expressed in Juv-s50 *∩* s56-s50 (Group B).

| **XI-PSID** | **Gene Symbol** | **Description** | **Juv-s56**<br>**Log<sub>2</sub> FC** | **s56-s50**<br>**Log<sub>2</sub> FC** |
| --- | --- | --- | --- | --- |
| XL24510.1.S1 at | dspp | dentin sialophosphoprotein | -8.9 | -6.3 |
| XL56635.2.S1 at | krt5.6 | keratin 5, gene 6 | -8.1 | -7.9 |
| XL24813.1.S1 at | krt14 | keratin 14, type I | -8.0 | -7.7 |
| XL9655.1.S1 at | neb | nebulin | -7.3 | -5.1 |
| XL51757.1.S1 at | umod.2 | uromodulin (uromucoid, Tamm-Horsfall glycoprotein), gene 2 | -7.1 | -6.1 |
| XL57075.1.S1 at | NA | NA | -7.0 | -6.7 |
| XL24625.1.S1 at | NA | NA | -7.0 | -7.0 |
| XL56635.1.S1 at | krt5.3 | keratin 5, gene 3 | -6.9 | -3.7 |
| XL14726.1.S1 at | NA | NA | -6.8 | -6.6 |
| XL11387.1.A1 at | hbz | hemoglobin, zeta | -6.4 | -6.4 |
| XL1321.2.S1 a at | NA | NA | -6.3 | -4.1 |
| XL25098.1.A1 at | NA | NA | -6.1 | -6.1 |
| XL28735.1.S1 at | tmem45b | Transmembrane protein 45B | -6.1 | -6.1 |
| XL31452.1.S1 at | myh1 | myosin, heavy chain 1, skeletal muscle, adult | -6.0 | -5.5 |
| XL11074.1.S1 at | tnni1.2 | troponin I type 1 (skeletal, slow), gene 2 | -5.9 | -5.7 |
| XL24517.1.A1 at | NA | NA | 6.0 | 6.7 |
| XL8551.1.S1 at | darmin | Darmin | 6.2 | 4.2 |
| XL9119.1.A1 at | NA | NA | 6.2 | 6.6 |
| XL9871.1.S1 at | thbs4 | thrombospondin 4 | 6.6 | 5.9 |
| XL48312.1.S1 at | slc13a5 | solute carrier family 13 (sodium-dependent citrate transporter), member 5 | 6.6 | 5.7 |
| XL9501.1.S1 at | nnmt | Nicotinamide N- methyltransferase | 6.6 | 5.6 |
| XL6687.1.S1 at | NA | NA | 6.8 | 4.6 |
| XL2009.2.S1 at | NA | NA | 6.9 | 6.7 |
| XL46955.1.S1 at | xpnpep2 | X-prolyl aminopeptidase (aminopeptidase P) 2, membrane-bound | 7.0 | 5.4 |
| XL46749.1.A1 at | NA | NA | 7.3 | 5.8 |
| XL17583.1.S1 at | krt5.2 | keratin 5, type II | 7.7 | 7.9 |
| XL17372.1.S1 at | lgalsia-a | galectin-Ia | 8.1 | 5.9 |
| XL19979.1.S1 at | bglap | Bone gamma-Carboxyglutamate (gla) protein | 8.5 | 7.8 |
| XL54855.1.S1 at | NA | NA | 8.7 | 8.0 |
| XL25162.1.A1 s at | NA | NA | 10.3 | 9.2 |

Xl-PSIDs that were differentially expressed in *s56*-*s50* ∩ *Juv*-*s56* comparisons (*s56_centric*) comprised the lowest count of DEX in Group B comparisons with a total of 98 (Fig. 3B), 49% of which were upregulated in *s56*. The *s56_centric* cohort of genes may represent differentially expressed genes whose expression peaks as metamorphosis approaches, and then decrease after its completion. This gene set could include genes with a role in preparing the inner ear for metamorphosis. In addition, the percentage of Xl-PSIDs without annotation was greater in this category than in *s56_centric* at 40%. Two of the top 15 Xl-PSIDs upregulated in *s56* were transcription factors (Table 6). The transcription factor annotated as specificity protein transcription factor (*sp7*) was among the most upregulated in *s56* with fold changes of −5.6 (*Juv*-*s56*) and 7.8 (*s56*-*s50*). A genetic mutation in *sp7* has been linked to osteogenesis imperfecta (Lapunzina et al., 2010). The other transcription factor in this category was the Xl-PSID annotated as distal-less homeobox 3 (*dlx3-b*), which was one of the lower expressed of the top 15 with fold changes of −4 (*Juv-s56*) and 3.2 (*s56-s50*). The three Xl-PSIDs with the greatest downregulation in *s56* were unannotated. The most downregulated annotated Xl-PSID was calcium binding and coiled-coil domain 1 (*calcoco1*) with fold changes of 4.9 (*Juv*-*s56*) and −3.4 (*s56-s50*). The *calcoco1* gene is considered a positive regulator of transcription and has been shown to be expressed in multiple tissues including the brain and kidney (Kim et al., 2003).

**Table 6.** Table of the top 15 up/downregulated Xl-PSIDs differentially expressed in Juv-s56 *∩* s56-s50 (Group B).

| **Xl-PSID** | **Gene Symbol** | **Description** | **Juv-s56**<br>**Log<sub>2</sub> FC** | **s56-s50**<br>**Log<sub>2</sub> FC** |
| --- | --- | --- | --- | --- |
| Xl.56532.1.S1 at | NA | NA | -6.2 | 6.3 |
| Xl.4801.1.S1 at | folr4 | Folate receptor 4 (delta) homolog (mouse) | -5.8 | 5.9 |
| Xl.8933.1.A1 at | sp7 | Sp7 transcription factor | -5.6 | 7.8 |
| Xl.17563.1.A1 x at | NA | NA | -5.6 | 5.5 |
| Xl.24484.1.A1 at | NA | NA | -5.5 | 5.6 |
| Xl.54837.1.S1 at | smpd3 | Sphingomyelin phosphodiesterase 3, neutral membrane(neutral sphingomyelinase II) | -5.1 | 3.9 |
| Xl.43319.1.S1 at | NA | NA | -5.0 | 4.9 |
| Xl.26062.1.S1 at | krt5.6 | keratin 5, gene 6 | -4.7 | 4.7 |
| Xl.24137.1.A1 at | NA | NA | -4.7 | 4.0 |
| Xl.1928.1.A1 at | NA | NA | -4.6 | 4.3 |
| Xl.55457.1.S1 at | pi15 | peptidase inhibitor 15 | -4.5 | 3.8 |
| Xl.52344.2.S1 s at | serpinh1 | serpin peptidase inhibitor, clade H (heat shock protein47), member 1, (collagen binding protein 1) | -4.4 | 4.6 |
| Xl.56226.1.S1 at | NA | NA | -4.3 | 4.0 |
| Xl.838.1.S1 at | dlx3-b | distal-less homeobox 3 | -4.1 | 3.2 |
| Xl.12285.1.S1 at | NA | NA | -4.0 | 4.1 |
| Xl.34628.1.A1 at | acaa2 | acetyl-CoA acyltransferase 2 | 2.7 | -2.2 |
| Xl.24287.1.A1 x at | rhov | ras homolog family member V | 2.7 | -3.5 |
| Xl.49447.1.S1 at | clip3 | CAP-GLY domain containing linker protein 3 | 2.8 | -1.6 |
| Xl.52118.1.S1 at | fam83a | family with sequence similarity 83, member A | 2.8 | -2.5 |
| Xl.51477.1.S1 at | NA | NA | 3.0 | -2.6 |
| Xl.1893.1.S1 at | hla-dqa1 | major histocompatibility complex, class II, DQ alpha 1 | 3.1 | -2.3 |
| Xl.2366.1.S1 at | pof1b | premature ovarian failure, 1B | 3.3 | -2.6 |
| Xl.10965.1.A1 at | NA | NA | 3.6 | -4.1 |
| Xl.28469.1.A1 at | umad1 | UBAP1-MVB12-associated (UMA) domain containing 1 | 3.8 | -2.8 |
| Xl.10626.1.A1 at | NA | NA | 3.8 | -2.8 |
| Xl.48800.1.S1 at | fntb | farnesyltransferase, CAAX box, beta | 3.9 | -5.5 |
| Xl.32224.1.S1 at | calcoco1-b | calcium binding and coiled-coil domain 1 | 4.9 | -3.4 |
| Xl.19817.1.A1 at | NA | NA | 6.6 | -6.9 |
| Xl.1996.1.A1 at | NA | NA | 6.9 | -4.8 |
| Xl.55124.1.S1 at | NA | NA | 9.5 | -8.5 |

The *juvenile centric* Xl-PSIDs, as shown in the Venn diagram in Fig. 3B, represented the most populated category with 432 Xl-PSIDs. This category had the fewest unannotated Xl-PSIDs at 33%, and 48% of the Xl-PSIDs were upregulated in the juvenile (Table 4). The two most upregulated Xl-PSIDs in juvenile were annotated as the same hemoglobin gene (*hba1*), with fold changes ranging from 8.7 to 11 (Table 7). Another Xl-PSID highly upregulated in juvenile was annotated as solute carrier family 25 [mitochondrial carrier; adenine nucleotide translocator], member 5 (*slc25a5*), with a fold change of 9 in both *Juv-s50* and *Juv-s56*.

**Table 7.** Table of the top 15 up/downregulated Xl-PSIDs differentially expressed in Juv-s50 *∩* Juv-s56 (Group B).

| **XI-PSID** | **Gene Symbol** | **Description** | **Juv-s50**<br>**Log<sub>2</sub> FC** | **Juv-s56**<br>**Log<sub>2</sub> FC** |
| --- | --- | --- | --- | --- |
| XL4816.1.S1 at | hba1 | hemoglobin, alpha 1 | 11.0 | 8.7 |
| XL17415.1.S1 at | hba1 | hemoglobin, alpha 1 | 10.9 | 9.1 |
| XL56128.1.S1 at | NA | NA | 10.0 | 10.0 |
| XL56717.1.S1 at | NA | NA | 10.0 | 10.1 |
| XL48636.1.S1 at | fmo3 | flavin containing monooxygenase 3 | 9.3 | 7.2 |
| XL53514.1.S1 at | slc25a5 | solute carrier family 25 (mitochondrial carrier; adenine nucleotide translocator), member 5 | 9.3 | 9.2 |
| XL24685.1.S1 at | NA | NA | 8.9 | 8.9 |
| XL9177.1.S1 at | NA | NA | 8.5 | 7.8 |
| XL6174.1.A1 at | rdd2 | NA | 8.3 | 8.3 |
| XL21584.1.S1 at | adh4 | alcohol dehydrogenase class IV | 7.8 | 6.3 |
| XL55367.1.A1 at | figf | c-fos induced growth factor (vascular endothelial growth factor D) | 7.6 | 5.6 |
| XL54279.1.S1 at | aoc3 | amine oxidase, copper containing 3 | 6.9 | 4.9 |
| XL47003.1.S1 at | mogat2.1 | monoacylglycerol O-acyltransferase 2, gene 1 | 6.6 | 6.6 |
| XL4790.1.A1 x at | NA | NA | 6.6 | 6.6 |
| XL9494.1.A1 at | pigr | polymeric immunoglobulin receptor | 6.5 | 6.3 |
| XL56666.1.S1 at | rhcg | Rh family, C glycoprotein | -7.4 | -8.0 |
| XL25492.4.S1 s at | ocm.2 | oncomodulin | -7.6 | -5.9 |
| XL1115.1.S1 at | actc1 | actin, alpha, cardiac muscle 1 | -7.9 | -5.2 |
| XL1125.1.S1 x at | hba3 | NA | -8.0 | -7.9 |
| XL54920.1.A1 x at | gstp1 | glutathione S-transferase pi 1 | -8.0 | -10.5 |
| XL7017.1.S1 at | krt12 | keratin 12, type I | -8.6 | -6.7 |
| XL54920.1.A1 at | gstp1 | glutathione S-transferase pi 1 | -8.9 | -11.5 |
| XL1126.1.S1 x at | hba4 | alpha-T4 globin, Putative ortholog of hemoglobin alpha chain [Source:Uniprot/SWISSPROT;Acc:P01922], 2 of 3 | -9.4 | -9.1 |
| XL1127.1.S1 at | hba2 | hemoglobin, alpha 2 | -9.7 | -9.3 |
| XL24530.2.A1 at | hbg2-b | Hemoglobin, gamma G | -10.3 | -10.2 |
| XL24530.1.S1 a at | hbg2-b | hemoglobin, gamma G | -10.4 | -10.4 |
| XL23992.1.S1 at | pla2g2e | phospholipase A2, group IIE | -10.7 | -9.2 |
| XL24530.2.S1 a at | hbg2-b | hemoglobin, gamma G | -11.1 | -11.0 |
| XL9653.1.S1 at | MGC53003 | parvalbumin beta-like | -11.3 | -9.9 |
| XL56832.1.A1 at | NA | NA | -12.1 | -9.1 |

Hemoglobin genes were also among the Xl-PSIDs with the greatest downregulation in juvenile. Twelve of the top 15 Xl-PSIDs downregulated in juvenile were annotated, six of which were annotated as hemoglobin genes. In addition, three of the annotated Xl-PSIDs were annotated as the same hemoglobin gene (*hbg2-b*), with fold changes ranging from −9.2 to −11.1.

### Analysis of Xl-PSIDs differentially expressed in all three comparisons identified genes with changing expression profiles throughout development (Group C)

This expression category contained the fewest DEX and contained the greatest proportion of unannotated Xl-PSIDs at 55%. There were equal percentages (41%) of DEX in all three stages with expression profiles that decreased or increased during development (Fig. 4 D-F). The annotated Xl-PSIDs that decreased during development were hemoglobin, epsilon 1 (*hbe1*); keratin 5, gene 6 (*krt5.6*); keratin 12 (*krt12*); aurora kinase b (*aurkb-b*); and protein kinase, cGMP-dependent, type II (*prkg2*) [Table 8]. The cohort of annotatedXl-PSIDs with increasing expression during development included dipeptidyl-peptidase 4 (*dpp4*); Corticotropin releasing hormone receptor 1, gene 2 (*crhr1.2*); and CD302 molecule (*cd302*). The other expression pattern observed had intensity values that increased from *s50* to *s56*, then decreased from *s56* to juvenile; 18% of the Xl-PSIDs possessed this pattern and the only one with a gene annotation was the presumed transcription factor Kruppel-like factor 5 (*klf5*). The gene *klf5* was also among the most differentially regulated genes in Group B, but was associated with a different Xl-PSID.

**Table 8.** Table of the 22 Xl-PSIDs differentially expressed in Juv-s50 *∩* Juv-s56 *∩* s56-s50 (Group C).

**8. Table of the 22 XI-PSIDs differentially expressed in Juv-s50 ∩ Juv-s56 ∩ s56-s50 (Group C).**
| **XI-PSID** | **Gene Symbol** | **Description** | **Juv-s50**<br>**Log<sub>2</sub> FC** | **Juv-s56**<br>**Log<sub>2</sub> FC** | **s56-s50**<br>**Log<sub>2</sub> FC** |
| --- | --- | --- | --- | --- | --- |
| Xl.57038.1.S1 at | ly6g6c | lymphocyte antigen 6 complex, similar to G6C | 9.8 | 4.8 | 5.0 |
| Xl.45569.2.S1 at | dpp4 | dipeptidyl-peptidase 4 | 8.1 | 5.5 | 2.7 |
| Xl.10081.1.A1 at | NA | NA | 7.8 | 3.9 | 3.9 |
| Xl.2904.1.S1 at | NA | NA | 7.7 | 5.5 | 2.2 |
| Xl.35284.1.S1 at | NA | NA | 7.4 | 3.9 | 3.5 |
| Xl.54254.1.S1 at | cd302 | CD302 molecule | 6.4 | 3.2 | 3.2 |
| Xl.12311.1.A1 at | NA | NA | 5.9 | 2.3 | 3.6 |
| Xl.52.1.S1 at | crhr1.2 | corticotropin releasing hormone receptor 1, gene 2 | 5.2 | 3.1 | 2.2 |
| Xl.42259.1.S1 at | NA | NA | 4.7 | -4.6 | 9.3 |
| Xl.55547.1.A1 at | klf5 | Kruppel-like factor 5 (intestinal) | 3.9 | -2.2 | 6.1 |
| Xl.30096.1.A1 at | NA | NA | 3.5 | 1.8 | 1.7 |
| Xl.26286.1.A1 at | NA | NA | 1.9 | -1.5 | 3.4 |
| Xl.28932.1.A1 at | NA | NA | -3.5 | -5.6 | 2.1 |
| Xl.4033.1.S1 at | aurkb-b | aurora kinase B | -3.6 | -1.7 | -1.9 |
| Xl.29732.1.S1 at | NA | NA | -4.7 | -2.2 | -2.5 |
| Xl.15081.1.A1 at | NA | NA | -4.7 | -3.1 | -1.5 |
| Xl.26004.1.S1 at | NA | NA | -4.8 | -2.4 | -2.4 |
| Xl.20138.1.A1 at | NA | NA | -5.2 | -2.7 | -2.5 |
| Xl.47670.1.S1 at | prkg2 | protein kinase, cGMP-dependent, type II | -6.7 | -3.4 | -3.3 |
| Xl.56635.2.S1 a at | krt5.6 | keratin 5, gene 6 | -7.6 | -3.8 | -3.9 |
| Xl.18637.1.S2 at | krt12 | keratin 12, type I | -10.9 | -9 | -1.9 |
| Xl.49005.1.S1 at | hbe1 | hemoglobin, epsilon 1 | -11.8 | -10.1 | -1.7 |

### DAVID analysis produced significant clusters from Group A and Group B comparisons

The Database for Annotation, Visualization and Integrated Discovery (DAVID) is a tool that can be used to uncover enriched gene clusters from lists of Xl-PSIDs. In addition to taking a gene-by-gene approach to identify genes with potential relevance to inner ear development and/or function, the use of tools such as DAVID can permit global analysis of many genes with similar characteristics, such as being upregulated in one developmental stage when compared to another.

DAVID analysis was undertaken using Xl-PSIDs lists from Groups A and B, which included six of the seven differential expression patterns identified in this study. The seventh differential expression pattern of differentially expressed in all pairwise comparisons (Group C) was excluded due to an insufficient number of Xl-PSIDs. To focus on stage specific functionality, the up- and downregulated Xl-PSIDs were analyzed separately. The Xl-PSID to DAVID ID mapping rate ranged from 33% to 57%, with most near or above the 50% mapping rate (Table 9). One member from Group A and two members from Group B generated significant clusters: 1) Xl-PSIDs upregulated in *s56* in the *Juv-s56* comparison; 2) Xl-PSIDs downregulated in *s50* in the *s50*_*centric* category (DEX in *Juv-s50* ∩ *s56-s50*); and Xl-PSIDs downregulated in juvenile in the *juvenile_centric* group (DEX in *Juv-s50* ∩*Juv-s56*). DAVID analysis of the Group A *Juv*-*s56* differential expression category produced a significant cluster from Xl-PSIDs upregulated in the *s56* stage. This cluster had an enrichment score of 1.6 (Benjamini adjusted p-value = 0.01) and it was associated with the term “*wnt signaling pathway*”. The *s50_centric* category from Group B produced a single significant cluster from Xl-PSIDs downregulated in *s50* (Benjamini adjusted p-value = 0.01) with an enrichment score of 1.8. The term associated with this cluster was “*calcium ion binding*”. Finally, the DAVID analysis generating both the greatest number of clusters and the clusters with the greatest significance were from Xl-PSIDs downregulated in juvenile in the Group B *juvenile_centric* differential expression category. The most enriched cluster had an enrichment score of 16.2 (Benjamini adjusted p-value = 2.02 x 10^-27^) and was composed of terms such as “cell cycle”, “cell division”, and “mitosis”. Other terms found in significant clusters included “microtubule motor activity”, “purine nucleoside binding”, “cyclin related”, and “oxygen transport”.

**Table 9.** Table summarizing the results of DAVID functional annotation.

|  | Group A |  |  |  |  |  | Group B |  |  |  |  |  |
| --- | --- | --- | --- | --- | --- | --- | --- | --- | --- | --- | --- | --- |
| | s56-s50 | | Juv-s56 | | Juv-s50 | | Juv-s50 ∩ s56-s50 | | Juv-s56 ∩ s56-s50 | | Juv-s50 ∩ Juv-s56 | |
|  | Up<br>s56 | Up<br>s50 | Up<br>Juv | Up<br>s56 | Up<br>Juv | Up<br>s50 | Up<br>s50 | Down<br>s50 | Up<br>s56 | Down<br>s56 | Up<br>Juv | Down<br>Juv |
| #Xl-PSIDs | 62 | 97 | 148 | 74 | 196 | 166 | 119 | 96 | 48 | 50 | 208 | 224 |
| #Mapped David IDs | 30 | 44 | 72 | 33 | 99 | 81 | 51 | 49 | 16 | 21 | 99 | 128 |
| % Mapped | 48% | 45% | 49% | 45% | 51% | 49% | 43% | 51% | 33% | 42% | 48% | 57% |
| Significant Clusters | 0 | 0 | 0 | 1 | 0 | 0 | 0 | 1 | 0 | 0 | 0 | 5 |

## Discussion

### Xl-PSIDs for genes encoding proteins with confirmed inner ear function were identified as differentially expressed

High-throughput technologies are powerful tools for transcriptomic analysis because they enable many genes to be profiled simultaneously, thereby facilitating the analysis of the relationships between genes with similar expression profiles. High-throughput technologies also provide a global perspective of gene expression in the system under investigation that can offer a rich context for focused studies of individual or smaller cadres of genes (Baldessari et al., 2005; McDermott et al. 2007; Darville and Sokolowski, 2013). The quality of the data is an essential component of experimental design that must be considered when generating and analyzing high-throughput data. Accordingly, we implemented a protocol to produce a microarray data set that was optimized to enhance reproducibility and minimize experimental variation generated by technical procedures. Evidence of this optimization can be seen in Fig. 1A, where the similarities in intensity distribution of the replicate data are apparent even prior to normalization.

Our microarray data analysis identified 1510 distinct DEX. The DEX included genes with established function in the inner ear, comprising genes associated with hearing loss and/or vestibular dysfunction such as transcription factors *sp7, and neurod1-b*. Neurod1 is expressed during otic vesicle specification and is known to be involved in both auditory and vestibular development (Liu et al., 2000; Alsina et al., 2009). Mutations in *neurod1* are linked to a clinical syndrome whose symptoms include sensorineural deafness (Rubio-Cabezas et al., 2010). In addition, genes were identified that are related to syndromic hearing loss such as structural proteins *col1a1* and *col1a2* (implicated in Osteogenesis imperfect [Marini et al., 2007]) and the transcription factor *tfap2a-b* (linked to Branchio-oculo-facial syndrome [Tekin et al., 2009]). The detection of known inner ear genes in this dataset enhanced confidence in the technique and demonstrated that transcripts detected in the inner ears of other vertebrates were also detected in the *Xenopus* inner ear.

### Pairwise comparisons uncovered Xl-PSIDs with seven differential expression patterns, including those for genes not previously associated with inner ear function

Pairwise comparisons between the three developmental stages produced seven differential expression patterns that were partitioned into Group A, B, or C contingent on whether an Xl-PSID was differentially expressed in one, two or three pairwise comparisons; respectively. Of these Xl-PSIDs, 64% corresponded to genes with an established functional annotation. Analysis of Group A Xl-PSIDs showed upregulation of gene expression in the *s50* and in the juvenile stages, and downregulation of expression in *s56*, with the greatest differences observed between s50 and juvenile stages (Fig. 4 A-C). Although the inner ears of the *s50* and juvenile animals are both functional, dramatic changes occur during this period such as an increase in the number of hair cells, axon projections, and in the size of the inner ear overall that are all likely contributors to the differences in gene expression observed (Díaz et al., 1995; López-Anaya et al., 1997; Serrano et al., 2001).

A concept implemented in this study is that of “*stage centricity*”, which designates sets of Xl-PSIDs in which the direction of expression in one stage (the centric stage) is in opposition to that of the other two stages. The comparisons that produced stage centric expression patterns were designated as Group B (Table 7, Fig. 3B). In Group B, the *s50_centric* Xl-PSIDs had the greatest proportion of upregulation with 55% upregulated in s50, indicating greater gene expression in the youngest stage when compared to both the s56 and the juvenile. The *juvenile_centric* expression category comprised 29% of the Group B Xl-PSIDs, twice as many as *s50_centric* and four times the number in *s56_centric*. This further supports the conclusion that a larger difference in gene expression is found between the juvenile and larval stages than between the two larval stages. In addition, there appears to be more downregulation of gene expression as the inner ear ages from larval to the juvenile stage.

The smallest cohort of Xl-PSIDs were found in Group C (Fig. 4 D-F, Table 8), which comprised the 22 Xl-PSIDs that were differentially expressed between all pairwise comparisons. Twelve of the Group C Xl-PSIDs did not have a gene annotation, making the most actively regulated Xl-PSIDs the least annotated category in this analysis. The dynamic nature of the genes that are changing expression throughout the developmental stages examined combined with the lack of annotation make these Xl-PSIDs of particular interest since their fluctuating expression levels may indicate a functional role in inner ear development and maturation that has yet to be characterized.

### Xenopus gene annotation limits interpretation of differential expression patterns

DAVID functional annotation clustering was undertaken on Xl-PSIDs from Groups A & B as summarized in Table 9. A total of seven significant clusters were discovered, five of which were found for Xl-PSIDs downregulated in juvenile in the *juvenile_centric* expression category. We noted that Xl-PSIDs downregulated in juvenile in the *juvenile_centric* set were both the most abundant (224 Xl-PSIDs), and had the greatest proportion of Xl-PSIDs that mapped to DAVID IDs (57%). A recurring theme encountered in this study was a lack of annotation for many of the differentially regulated genes. Three of the seven differential expression categories that produced significant clusters included terms previously associated with inner ear function or development, but the inability to map many of the Xl-PSIDs to DAVID IDs likely had a negative impact on the effectiveness of the functional analysis.

### Toward a Xenopus model for hearing and balance

The inner ear remains a relatively understudied organ. Data from this *Xenopus* transcriptome-wide study can be mined to identify new candidate genes for inner ear function and to explore unrecognized relationships between inner ear genes. The dataset presented here complements and extends genetic outcomes from inner ear research that has focused on induction and formation of the otic placode (Giraldez and Fritzsch, 2007; Alsina et al, 2009; Almasoudi and Schlosser, 2021).. The observation that approximately a third of the differentially expressed *Xenopus* inner ear genes have no annotation highlights the prevalence of knowledge gaps regarding genes that are involved in sensory organ development. These unannotated genes represent an exciting opportunity to fully characterize the genetics underlying inner ear development and to identify genes crucial for balance and audition. Additionally, when genetic characterizations are carried out in an amphibian like *Xenopus*, we have the future prospect of identifying genes involved in hair cell regeneration, a process that does not occur easily in the mammalian inner ear (Costa et al., 2015; Lee and Waldhaus, 2022). It is therefore anticipated that the data set presented here will become a resource that can be used to generate hypotheses regarding genes and mechanisms that underlie inner ear development and repair.

In summary, *Xenopus laevis* is an established model for vertebrate development and cellular biology, especially during embryogenesis (reviewed by Harland and Grainger, 2011; Gurdon, 2014), but its allotetraploid genome historically was not easily amenable to genetic manipulation. Previously this limitation was overcome using the related diploid species *X. tropicalis*, however; the introduction of methods such as transcription activator-like effector nuclease (TALEN) and CRISPR-Cas have made it possible to manipulate the *X. laevis* genome directly (Nakade et al., 2015; Naert et al, 2017; Horb et al. 2019). Moreover, the sequencing of genomes from both *X. tropicalis and X. laevis* (Hellsten et al., 2010; Exner and Willsey, 2020) has enhanced the value of *Xenopus* in the analysis of vertebrate organ development and gene function. These features, in conjunction with the current data set, afford advantages to *Xenopus* as a model organism for biomedical investigations of the development of the senses of hearing and balance.

## Materials and Methods

### Xenopus laevis

*Xenopus laevis* larvae and juvenile [*Juv*] animals were purchased from Nasco (Fort Atkinson, WI). The youngest larval age group represented in this study was comprised of animals staged 50 thru 52 (Avg. Wt. 0.15 g ± 0.03 g), designated as *s50*; the other larval age was comprised of animals staged 56 thru 58 (Avg. Wt. 0.54 g ± 0.1 g); designated as *s56*. The juvenile stage consisted entirely of post metamorphic animals (Avg. Wt. 9.43 g ± 0.3 g; 3-5 months of age). The New Mexico State University Institutional Animal Care and Use Committee (IACUC) approved all animal procedures. Staging of animals was based on anatomical landmarks determined by Nieuwkoop and Faber (Nieuwkoop and Faber, 1967).

### RNA Isolation and replicate sample preparation

*Xenopus* inner ears were dissected as previously described (Trujillo-Provencio et al, 2009). Dissected inner ear tissue was removed from RNAlater and homogenized with a Brinkman Polytron PT1200. Total RNA was extracted following manufacturer’s directions supplied with the RNeasy^®^ Mini Kit (Qiagen). Total RNA was purified using the DNA-*free* kit^TM^ (Ambion).

A single replicate consisted of RNA extracted from the inner ears of multiple animals. Three replicates were produced for each stage, for a total of nine unique samples. The number of animals used for each replicate varied by stage due to differences in inner ear size and weight (*s50*: 15, 38, 24; *s56*: 10, 7, 8; juvenile: 5, 5, 6). RNA from each replicate was quantified and assessed for quality using the Agilent Technologies 2100 Bioanalyzer. All replicates used for microarray analysis scored a minimum RNA integrity number (RIN) value of 9.0.

### Sample amplification and microarray hybridization

*Xenopus* inner ear RNA was profiled using the *Affymetrix* GeneChip^®^ *Xenopus laevis* Genome 2.0 Array (*Affymetrix*) comprising more than 32,400 probe sets (Xl-PSIDs), representing over 29,900 *X. laevis* transcripts. Labeled antisense single-stranded cDNA was prepared from each RNA replicate using the Ovation™ RNA Amplification System V2 in conjunction with the FL-Ovation™ cDNA Biotin Module V2 (both from NuGEN), hybridized overnight and scanned using a GeneChip Scanner 3000 7G (*Affymetrix*). RNA and microarrays were processed at the MIT BioMicro Center.

### Preprocessing for microarray Analysis

The original (raw) data from *X. laevis* GeneChip^®^ CEL files acquired from three replicate arrays per developmental stage were preprocessed using the Gene Chip robust multichip averaging (GCRMA) method to produce a single log_2_ transformed measure of the intensity level (in arbitrary units [a.u.] of fluorescence) for every Xl-PSID on each replicate array. The JMP Genomics 5.0 analysis platform was used for both preprocessing and differential expression analysis. The preprocessed microarray data served as the starting point for all downstream analysis. Data can be accessed at the Gene Expression Omnibus under accession numbers GSE69546, GSE73828, GSE73829.

### Differential expression analysis and pairwise comparisons

Pairwise comparisons were made between all stages using Analysis of Variance (ANOVA) as implemented by the JMP Genomics 5.0 analysis platform. Candidate Xl-PSIDs for differential expression were detected by analyzing pairwise differences in the average fluorescence intensity of the replicate Xl-PSIDs for each stage. Analysis of pairwise comparisons was undertaken for three groups comprising seven differential expression categories: Group A) differential expression in one pairwise comparison (*s56*-*s50*; *Juv*-*s56*; *Juv*-*s50*); Group B) differential expression in two pairwise comparisons (*Juv-s50* ∩ *s56-s50*; *Juv-s56* ∩ *s56-s50*; *Juv-s56* ∩ *Juv-s50*); Group C) differential expression in three pairwise comparisons *s56-s50* ∩ *Juv-s56* ∩ *Juv-s50*). A fourth possible group, “*different in none*” was not examined because the objective of this study was to uncover differentially expressed genes.

Filters and fold change restrictions were applied to the preprocessed data to identify candidate genes for differential expression during development. A q-value (≤ 0.01) established criteria for a positive false discovery rate (pFDR). The minimum fold change value for pairwise comparison was set to ±1.5 to allow for the capture of subtle changes in expression and to filter out potentially insignificant changes. As per Powers et al. (2012), only Xl-PSIDs that met the additional minimum average normalized intensity criteria of fluorescence intensity ≥ 4 A.U. on the replicate arrays were designated “upregulated”.

The resulting data were exported as a tab-delimited text file for the application of significance filters. A negative fold change in a given pairwise comparison was used to identify downregulated Xl-PSIDs. Java software was developed to apply the filters against the JMP output file. Following application of the fold change and fluorescence intensity significance filters, the differentially expressed data were stored in a MySQL database table. The Java software developed for this project will be made available by the authors and deposited to the GitHub open source project repository (https://github.com/).

### Annotation of Xl-PSIDs

Xl-PSIDs were annotated using a two-stage approach. Data files were downloaded from the *Xenbase* website (www.xenbase.org; Bowes et al., 2010) and formatted for insertion into a local MySQL database to facilitate automated queries. Differentially expressed Xl-PSIDs were linked to *Xenbase* annotation by searching data originating from the *Xenbase* data file GenePageAffymetrix_laevis2.0.txt (file date stamped January 29, 2015) which associated *Affymetrix* Xl-PSIDs to *Xenopus* gene symbols. Xl-PSIDs that remained unannotated by Xenbase were subjected to a second round of annotation process. The representative public id from the *Affymetrix* annotation file, X_*laevis*_2.na33.annot.csv, was queried against the *Xenopus laevis* UniGene build 94 Xl.data file for retrieval of informative annotation. Gene descriptions were obtained by linking *Xenbase* XB-Gene ID to data obtained from the files GenePageEnsemblModelMapping.txt or XenbaseGeneHumanOrthologMapping.txt (both files date stamped January 29, 2015), genes for which descriptive information was unavailable from these two files were subjected to a manual search on the NCBI Gene website at the following URL: http://www.ncbi.nlm.nih.gov/gene/. Xl-PSIDs still unannotated after the annotation approach described above were assigned “NA” as gene symbol. In addition, gene symbols derived from the Mammalian Gene Collection (http://mgc.nci.nih.gov) and those with a generic locus assignment were designated as unannotated unless associated with a specific gene description. Gene annotations were used to impart a functional connotation to the differentially expressed Xl-PSIDs identified in the pairwise comparisons. When associating Xl-PSIDs to gene symbols, the convention employed by *Xenbase* was adopted.

### Global analysis of pairwise comparison data

The Database for Annotation, Visualization and Integrated Discovery (DAVID) functional annotation clustering analysis was used to analyze the Xl-PSID lists generated for each pairwise comparison. The Xl-PSID lists were uploaded to the DAVID website and the Affymetrix chip version was selected for DAVID. The data lists were functionally annotated by selecting the analysis method “*Functional Annotation Clustering*” and the cluster analysis categories “SP_PIR_KEYWORDS” and “GO_TERM_MF_FAT” (to select the most specific GO terms). Upregulated and downregulated Xl-PSIDs were analyzed separately for each pairwise comparison using a clustering stringency of “Medium”. Clusters (and associated terms) with Benjamini adjusted p-values ≤ 0.05 were considered significant. Only clusters and terms meeting this criterion were reported as “enriched”.

## Acknowledgements

The authors wish to thank Dr. TuShun R. Powers for scientific advice in preparation of the manuscript and the MIT BioMicro Center for all aspects of microarray processing. The research reported in this publication was supported by the National Institute of General Medical Sciences of the National Institutes of Health under award numbers R25GM061222 and P50GM68762. The content is solely the responsibility of the authors and does not necessarily represent the official views of the National Institutes of Health

## Notes

https://www.ncbi.nlm.nih.gov/geo/query/acc.cgi?acc=GSE69546

https://www.ncbi.nlm.nih.gov/geo/query/acc.cgi?acc=GSE73828

https://www.ncbi.nlm.nih.gov/geo/query/acc.cgi?acc=GSE73829

### Competing Interest Statement

The authors have declared no competing interest.

